# Restoring ancestral phenotypes by reduction of plasticity is a general pattern in gene expression during adaptation to different stressors in *Tribolium castaneum*

**DOI:** 10.1101/683771

**Authors:** Eva L. Koch, Frédéric Guillaume

**Author notes:** Corresponding author: Prof. Dr. Frédéric Guillaume.

## Abstract

Plasticity and evolution are two processes individuals to respond to environmental change, but how both are related and impact each other is still controversial. We studied plastic and evolutionary responses in gene expression of *Tribolium castaneum* after exposure to new environments that differed from ancestral conditions in temperature, humidity or both. Using experimental evolution with ten replicated lines per condition, we were able to demonstrate adaptation after 20 generations. We measured gene expression in each condition in adapted selection lines and control lines to infer evolutionary and plastic changes. We found more evidence for changes in mean expression (shift in the intercept of reaction norms) in adapted lines than for changes in plasticity (shifts in slopes). Plasticity was mainly preserved and was responsible for a large part of the phenotypic divergence in expression between ancestral and new conditions. However, we found that genes with the largest evolutionary changes in expression also evolved reduced plasticity and often showed expression levels closer to the ancestral stage. Results obtained in the three different conditions were similar suggesting that restoration of ancestral expression levels during adaptation is a general evolutionary pattern. We increased the sample size in the most stressful condition and were then able to detect a positive correlation between proportion of genes with reversion of the ancestral plastic response and mean fitness per selection line.

## Introduction

Whenever facing environmental change, populations can adapt to new phenotypic optima by plasticity and evolution. Plasticity is the ability of a single genotype to produce multiple phenotypes as a function of the environment. It is often seen as an immediate response of individuals to changes in their environment. In contrast, evolution requires a change in allele frequencies within a population. This process occurs over several generations and represents a more long-term response, which can result in local adaptation. It is still not well understood how these two processes are related and interact with each other (de Jong, 2005; Forsman, 2015; Ghalambor, McKay, Carroll, & Reznick, 2007; Price, Qvarnström, & Irwin, 2003; Wund, 2012).

Plasticity does not affect a population’s genetic composition and should thus not have long-term consequences in the following generations. However, it changes the distribution of phenotypes on which selection can act (de Jong, 2005; Pfennig et al., 2010; Pigliucci, 2005; Price, Qvarnstrom, & Irwin, 2003). If plasticity perfectly matches the new phenotypic optimum, it prevents evolution since no genetic changes are required and selection is prevented (Ghalambor et al., 2007). On the other hand, plasticity is also crucial for a population’s persistence and can reduce the costs of selection (Chevin, Lande, & Mace, 2010; Pavey, Collin, Nosil, & Rogers, 2010). It prevents extinction and protects populations from bottleneck effects, thereby maintaining a higher genetic variation on which subsequently selection can act (Fitzpatrick, 2012; Pfennig et al., 2010; Massimo Pigliucci, 2005). There is both, theoretical (Chevin et al., 2010; Draghi & Whitlock, 2012; Fierst, 2011) and empirical work (Schaum, Rost, Millar, & Collins, 2013), demonstrating that more plastic populations exhibit faster evolution. The benefits of plasticity for persisting in new habitats were also demonstrated in invasive species (Molina-Montenegro, Peñuelas, Munné-Bosch, & Sardans, 2012; Pichancourt & van Klinken, 2012; Yeh & Price, 2004).

The extent of plasticity can be represented as a reaction norm (Scheiner, 1993), which is the phenotypic trait value as a function of an environmental variable. Evolution can affect the reaction norm in two ways: The intercept can be shifted corresponding to a change in the mean phenotypic value. Alternatively, the slope of the reaction norm, i.e. the plasticity, can be changed. Thus, plasticity itself can also be subject to evolution given that there is sufficient genetic variation in reaction norms (Garland & Kelly, 2008; Nussey, Postma, Gienapp, & Visser, 2005).

Dependent on whether plastic responses are adaptive and increase fitness of an individual, different patterns describing the relationship between ancestral plasticity and evolution are possible. If plastic responses are adaptive, but not sufficient to reach the phenotypic optimum, evolution should work in the same direction as the plastic response (referred to as Baldwin effect (Crispo, 2007) or cogradient variation (Conover, Duffy, & Hice, 2009). In this case, selection may favour the most plastic individuals, causing evolved populations to exhibit a higher plasticity than their ancestors (Crispo 2007; Lande 2009). Another possible outcome is genetic assimilation: An initially environmentally induced phenotype can get fixed by a loss of plasticity and becomes continuously expressed even in the ancestral environment (Levis & Pfennig, 2016; Pigliucci, Murren, & Schlichting, 2006). In contrast, if plastic responses are maladaptive, we expect to observe evolutionary changes opposite to plasticity (countergradient variation ((Conover et al., 2009) or genetic compensation (Grether 2005)). Maladaptive plasticity was proposed as a possible mechanism promoting evolution since it moves phenotypes further away from the optimum and thereby increases the strength of selection on the phenotype (Ghalambor et al., 2007). Both co-gradient (Barton, Sunnucks, Norgate, Murray, & Kearney, 2014; Conover et al., 2009) and counter-gradient (Conover et al., 2009; Ghalambor et al., 2015; Laugen, Laurila, Räsänen, & Merilä, 2003) evolutionary changes have been found, indicating that adaptive and maladaptive plasticity are common. Reversion of ancestral plasticity occurs more frequently (Ho & Zhang, 2018), indicating that plastic responses are often not beneficial for long-term adaptation.

Plastic responses in physiology, behaviour or morphological traits are often initiated by changes in gene expression (Hodgins-Davis & Townsend, 2009; Wray, 2007). The transcriptome represents a direct link between genotype and phenotype making it particularly interesting to study the interplay between plasticity and evolution. Transcription is highly plastic and modulating expression levels is an important part of an organism’s physiological adjustment to environmental change (Gibson, 2008; McCairns & Bernatchez, 2009). On the other hand, there are also many studies demonstrating evolutionary divergence in gene expression between locally adapted populations (Alvarez, Schrey, & Richards, 2015; Guo et al., 2016; Romero, Ruvinsky, & Gilad, 2012; Townsend, Cavalieri, & Hartl, 2003; Whitehead & Crawford, 2006). Gene expression may even evolve more rapidly than changes in proteins since mutations affecting the magnitude of expression are less likely to be deleterious than changes in protein structures (Carroll, 2005; Wray, 2007). However, it is not clear how fast plasticity in gene expression can change. Some studies reported changes in plasticity in few genes after adaptation to new conditions (Morris et al., 2014; Passow et al., 2017; von Heckel, Stephan, & Hutter, 2016), whereas others found only limited evolution of plasticity (Yampolsky, Glazko, & Fry, 2012) or less than expected (Huang & Agrawal, 2016).

In our study, we used whole transcriptomes to understand the interplay between plasticity and evolution at the gene expression level during adaptation to new environments with the model organism *Tribolium castaneum* (red flour beetle) using an experimental evolution approach. With RNA-seq, we quantified plastic and evolutionary changes in gene expression and their contribution to the total phenotypic divergence between populations inhabiting different environments. More specifically, we were interested to test whether the same genes exhibited both evolutionary and plastic changes in a new environment and whether evolved changes were in the same direction as their ancestral plasticity. In doing so, we could test how plasticity affected evolution. We tested whether plasticity prevented evolution, or whether maladaptive plasticity increased the strength of selection and promoted evolution. Next, we tested how evolution affected gene expression reaction norms, i.e. whether adaptation to new conditions was achieved by changes in slopes or intercepts.

## Material and Methods

### Animal rearing, experimental evolution

We used the *Tribolium castaneum* Cro1 strain (Milutinović, Stolpe, Peuß, Armitage, & Kurtz, 2013), collected from a wild population in 2010 and adapted to lab standard conditions (32°C, 70% relative humidity (r.h.)) for more than 20 generations. Beetles were kept in 24h darkness on organic wheat flour mixed with 10% organic baker’s yeast. We sterilized flour and yeast by heating them for 12h at 80°C before use. To test for adaptation to new environmental conditions we used replicate lines and exposed them to three treatment and control (CT) conditions. The conditions in the treatments were: Dry (D): 33°C and 30% r. h.; Hot (H): 37°C and 70% r. h.; Hot-Dry (HD): 37°C and 30% r. h. To generate replicate lines, we used 120 individuals (60 females and 60 males in the pupal stage) and placed them into a vial containing 80g medium. We produced six lines per selection regime (treatments plus control), resulting in a total of 24 lines. For each new generation, we randomly collected 120 pupae and placed them into a new vial. After seven to ten days, in which the pupae became adults, mated and laid eggs, adult beetles were removed by sieving the medium. We waited until the next generation (eggs/larvae in the medium) had reached the pupal stage and again collected 120 pupae per line to establish the next generation. This is similar to natural selection since individuals, depending on their fitness, do not contribute equally to the next generation. In generation 15 we produced additional mixed lines to prevent loss of genetic diversity by gene drift and inbreeding, which might impede adaptation: We mixed the six replicate lines of each selection regime in equal proportions (20 individuals from each replicate line) four times, resulting in four mixed lines with 120 individuals each. In total we had 39 lines: six normal and four mixed lines per selection regime (one line in D became extinct). The transplant experiment to test for adaptation was conducted in generation 22.

### Reciprocal transplant and fitness assay

Before testing for adaptation, all lines stayed for two generations in the same condition to remove potential maternal or epigenetic effects (Supporting information, Figure S1): Beetles of generation 20 from all selection lines were transferred to control conditions, in which they stayed for one week to mate and lay eggs. After removal of the adults, we waited until their offspring had reached the pupal stage and separated males and females. These individuals (generation 21) developed completely in control conditions. When they reached the adult stage, we created 13 full-sib families per selection line by transferring one virgin male with one virgin female of the same selection line in 15mL tubes with 1g of medium. After four days, in which the beetles could mate and lay eggs, 9g of medium was added to provide food for the developing offspring and each mating pair was transferred to a new vial. We repeated this three times, resulting in four vials per mating pair containing medium and eggs. Immediately after removal of the mating pair, vials of each mating pair were randomly assigned to the four different conditions, resulting in full-sib families split across all conditions. These beetles were transferred to the treatments at the egg stage. As soon as offspring in these vials had reached the pupal stage, males and females (four females and four males per family and condition) were separated and transferred to 15 mL tubes with 5 g of medium and remained there until they were used for the fitness assay two weeks later. They developed completely in treatment conditions. We then assessed their performance in each condition by estimating their fitness to test for adaptation. A virgin male and a virgin female of the same selection line from the same condition but from different families were again placed into a 15 mL tube with 1 g medium. After four days, the mating pair was removed. Males and females were transferred to 1 mL Eppendorf tubes (one individual per tube), immediately frozen in liquid nitrogen and stored at −80°C to use them for gene expression measurements. 9 g medium was added to the mating tube. After four weeks (in CT and H) or five weeks (D and HD), all offspring had reached the adult stage and were counted. We used the number of adult offspring as an estimate of the fitness of a mating pair.

### Statistical analysis

To test whether selection regime significantly influenced number of offspring produced and test whether 20 generations in the treatments resulted in adaptation, we compared offspring numbers of selection lines in their native condition to CT-lines transferred to the same condition. We applied linear mixed models using the R-packages *lme4* (Bates, Mächler, Bolker, & Walker, 2015), and *lmertest* (Kuznetsova, Brockhoff, & Christensen, 2017) and *lsmeans* (Lenth, 2016) to obtain *p*-values and confidence intervals. We included line and family as random factors, selection and line type (mixed/normal) and their interaction as fixed effects. To test whether the selection regime influenced how lines responded to the treatments, we used a linear mixed model with offspring number in CT and treatment conditions as response variable, condition, selection regime, line type (normal/mixed) and interactions as fixed effects and line, family and interaction between line and condition as random effects. A significant interaction between condition and selection regime indicates a significant effect of the selection regime (evolution) on the response to the conditions (plastic response).

### RNA extraction, library preparation and sequencing

208 female beetles (Table 1) p stored at −80°C were homogenized in Tri-Reagent® (Zymo Research, California, USA) using an electric bead mill. RNA was extracted with the RNA Mini Prep kit (Zimo Research, California, USA) following the instructions of the manufacturer. RNA-quality was checked on a TapeStation (Agilent, Waldbronn, Germany) and concentrations were measured with aQubit® Fluorometer (Life Technologies, California, USA). Libraries were created with 500 ng RNA for each individual separately with the LEXOGEN mRNA-Seq Library Kit following the manual (LEXOGEN GmbH, Vienne, Austria). Library quality was checked on a TapeStation (Agilent, Waldbronn, Germany) and concentrations were determined by qPCR. Libraries were diluted to the same molarity and pooled (33-36 libraries per pool). All treatments and selection regimes were randomized during RNA-extraction, library preparation, and sequencing. Single-end sequencing was performed in five runs on the Illumina NextSeq 500 (Illumina, Inc, California, USA) using the 75 cycles High Output Kit. After quality control using FastQC (www.bioinformatics.bbsrc.ac.uk/projects/fastqc) reads (adaptors were trimmed and the first 10 bases were hard trimmed, minimum average quality Q10, minimum tail quality 10, minimum read length 20) were mapped against the reference genome (ftp://ftp.ensemblgenomes.org/pub/release30/metazoa/gtf/tribolium_castaneum/Tribolium_castaneum.Tcas3.30.gtf.gz) with STAR v.2.5 (Dobin et al., 2013). We then used FeatureCounts (Liao, Smyth, & Shi, 2014) to count the number of reads that mapped to each gene in the reference genome. Mapping as well as read counting was performed within the data analysis framework SUSHI (Hatakeyama et al., 2016).

**Table 1:**
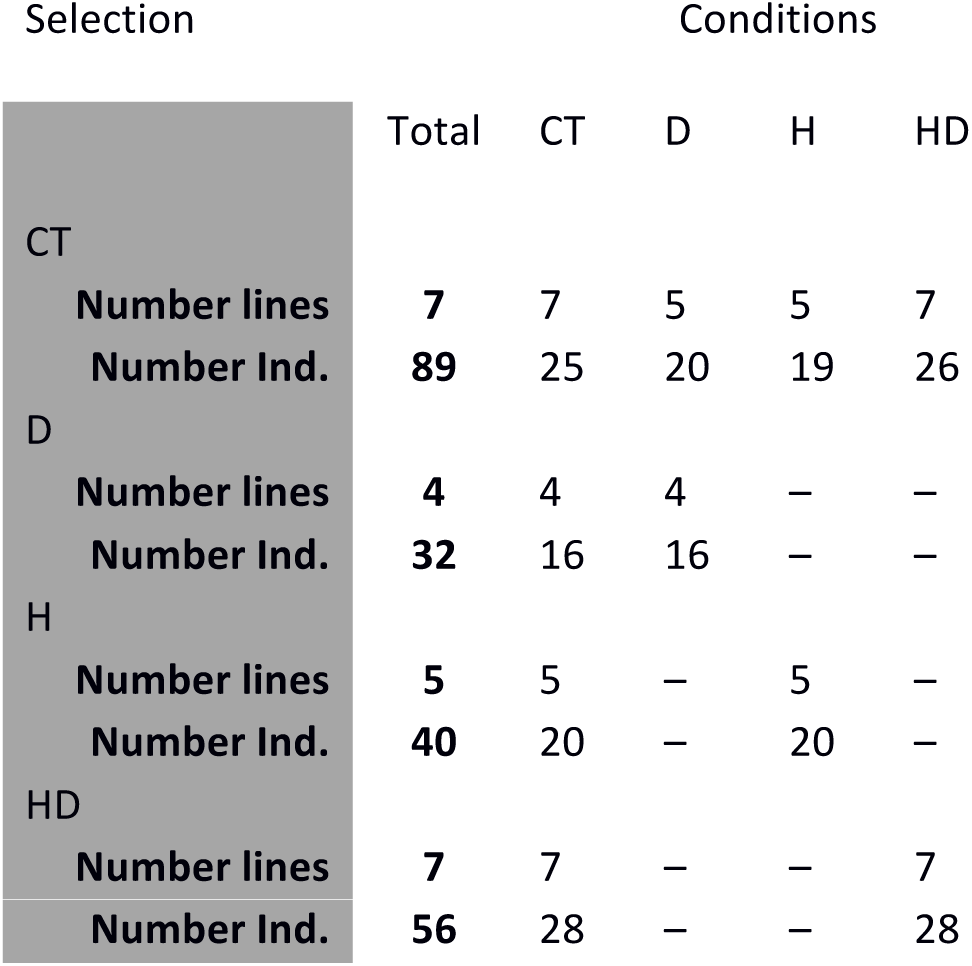
Number of sequenced replicate lines and individuals per selection and treatment, which were used for this study. Selection lines could adapt to conditions for 20 generations. Control (CT) conditions: 33°C, 70% relative humidity (r.h.); treatments: Dry (D): 33°C, 30% r.h.; Hot (H): 37°C, 70% r.h.; Hot-Dry (HD): 37°C, 30% r.h.

### Gene expression analysis

Gene expression analysis was done in R (R Core Team, 2017). We used the R package *edgeR* (Robinson, McCarthy, & Smyth, 2010) for normalizing (method: TMM) expression data to cpm (counts per million) after filtering lowly expressed genes (minimum of one cpm in at least two samples). For subsequent differential expression analysis we used the R package *limma* (Law, Chen, Shi, & Smyth, 2014; Ritchie et al., 2015). From the differential expression analysis, we obtained the number of differentially expressed genes (DE genes) within lines between conditions (plastic changes, see Figure S1) or between lines of different origins (CT vs. selection) within conditions (evolutionary changes, see Figure S1). The total phenotypic divergence in gene expression between CT and treatments (i.e. total change TC) is the differential expression (log2-fold change) between CT-lines in CT and selection lines in the treatments (Figure 1A-C and Supporting information Figure S1). The ancestral plasticity (PC_CT_) is the differential expression of CT-lines between CT and treatment conditions, while the evolved plasticity (PC_Sel_) is the same difference measured in selection lines. The evolutionary changes are EC_T_ when measured as differential expression between CT and selection lines in the treatments and EC_CT_ when measured in CT (Figure 1). Finally, differences between plastic responses of CT- and selection lines (the interaction between condition and selection regime) give the evolutionary change in plasticity. To partition TC into changes explained by ancestral plasticity (PC_CT_) and evolutionary changes (EC_T_, see Figure 1A-C), we calculated the relative contribution of each component to the total. We used the log2-fold change of each transcript to evaluate and compare the magnitude of the plastic and evolutionary changes. When computing log2-fold changes, we accounted for non-independence among individuals from the same line by using the *duplicateCorrelation* function (Smyth et al. 2005) and added sequencing runs as batch effect. A gene is classified as differentially expressed (DE) with a FDR ≤ 5% after adjusting for multiple testing (Benjamini and Hochberg 1995).

**Figure 1:**
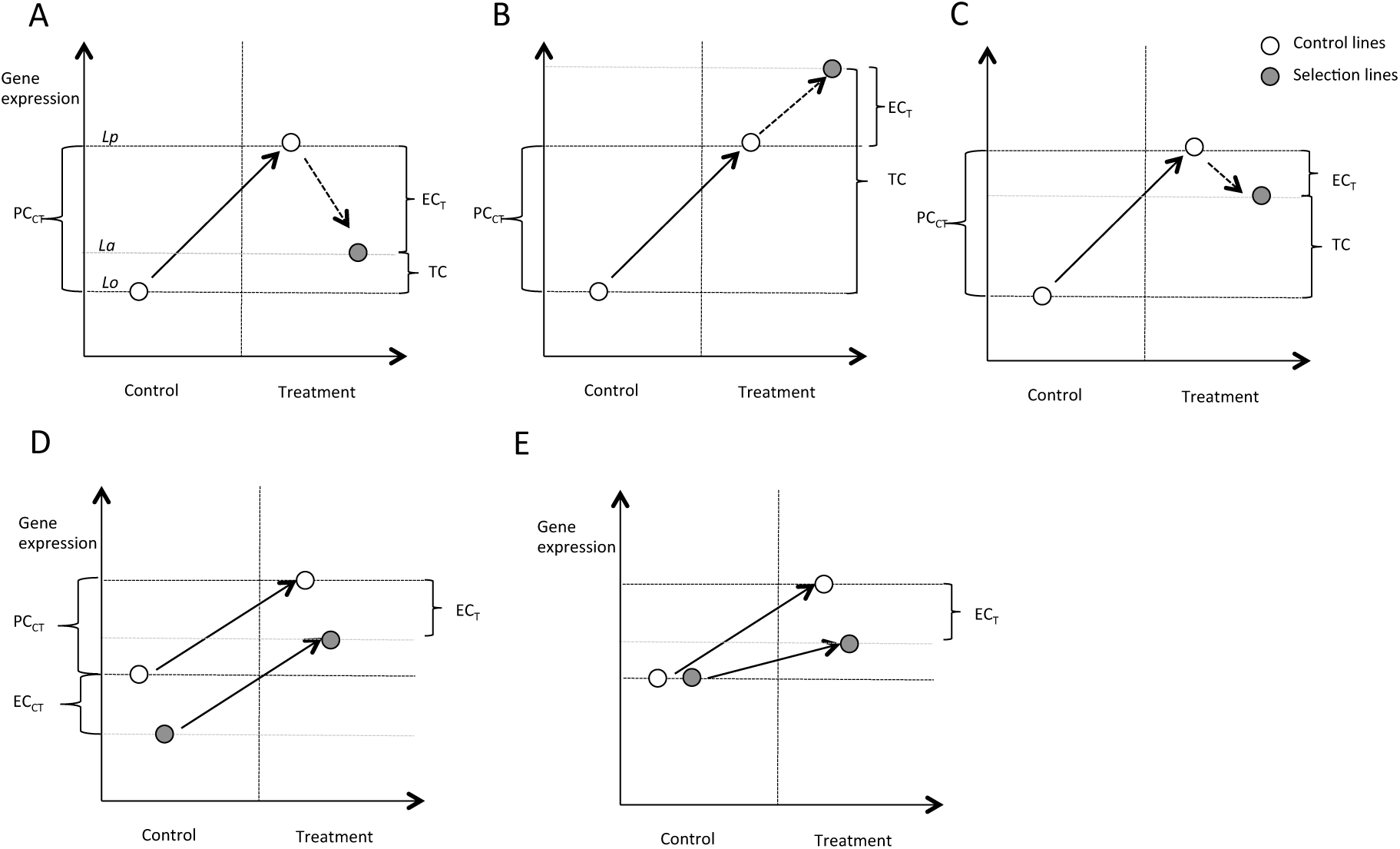
Possible relations between plastic change PC and evolutionary change EC. Gene expression levels of control lines (white) in the ancestral control condition represent the original stage Lo, expression in the treatment the plastic stage Lp. Expression levels of selection lines (grey) in the treatment give the adapted stage La Arrows indicate the direction of PC (dashed line) and EC (solid line). PC can be in opposite direction to EC (reversion **(A))** or it can be in the same direction (reinforcement **(B))**. However, even if PC and EC are opposite to each other, PC can bring expression levels closer to levels of the adapted lines. In this case, the total change TC, (difference between expression levels of control lines in control conditions and selection lines in treatment) is larger than EC **(C)**. During adaptation, lines could have reached the optimum by either changing mean expression, i.e. shift in the intercept of the reaction norm **(D)** or by changing their plasticity, i.e. the slope of the reaction norm **(E)**. In case of a change in the mean, plastic changes of Control lines PC_CT_ and selection lines PC_Sel_ as well as observed evolutionary change EC_T_ in treatment and in Control EC_CT_ would be highly correlated because reaction norms (red arrows) remain parallel **(D)**. If observed EC_T_ in treatment is due to a change in plasticity only, no correlation between EC_T_ and EC_CT_ should exist, and the correlation between PC_Sel_ and PC_CT_ should be small **(E)**.

In a second step, we used the normalized read counts (cpm, TMM-normalized) corrected for batch effects (sequencing runs) using the *removeBatchEffect* function in the *limma* R package (Ritchie et al., 2015). We used these counts to quantify expression levels in in CT and treatments. This allowed us to conduct an analysis for each selection lines separately as was described by Ho & Zhang (2018) (details see below).

### Evolution of reaction norms

Reaching a new phenotypic optimum can be achieved by changing the mean (intercept of the reaction norm) or by changing the plasticity of a trait (slope of reaction norm) (Figure 1D,E). We were interested in assessing which of these patterns was more prevalent in gene expression evolution. To test for an evolutionary change in plasticity (slope), we first evaluated the similarity in plasticity between CT- and selection lines by calculating the correlation between PC_CT_ and PC_Sel_ (Suppurting information Figure S2A) among all genes, independent of the significance. Second, to test for an evolutionary change in mean expression (intercept), we calculated the correlation between the evolved differences in CT (EC_CT_) and in treatment conditions (EC_T_). A significant correlation indicates that the overall mean has changed (Figure S2A). Further, for all genes with significant EC_T_, we quantified how much of this difference could be explained by a shift in the mean or by a change in plasticity. A shift in the mean corresponds to EC_CT_, whereas the remaining difference (EC_T_-EC_CT_) gives the evolutionary changes in plasticity (see also Stoks et al. 2016). Relative contributions of each component were obtained by dividing them by the total EC_T_.

To test for significant effects of the selection regime on differential expression (i.e., on number of DE genes, log2-fold change, and correlations), we used a permutation test. We randomly assigned samples and their transcriptomes to either CT selection or treatment (D, H, HD) selection (number of samples for each selection was not changed) and repeated the DE analysis. We kept the original assignment to lines and conditions and repeated the DE analysis for each permuted data set. Observed values (e.g. number of DE genes, correlations) were considered significant if higher than the most extreme 5% of the distribution calculated from permutations.

### Comparing plastic and evolutionary responses

To infer the relationship between ancestral plasticity and evolution, we compared the direction of PC_CT_ to the direction of EC_T_. Evolution may *reinforce* the plastic response when PC_CT_ is in the same direction as EC_T_ (Figure 1B). If EC_T_ is in opposite direction, it *reverses* PC_CT_ (Figure 1A). To test which of these patterns was more prevalent, we followed Ho and Zhang 2018 (Ho & Zhang, 2018). Expression levels (cpm) of CT-Lines in CT conditions represented the original stage (*Lo*), CT-Lines in treatment the plastic stage (*Lp*), and selection lines in their respective condition the adaptive stage (*La*) (Figure 2). For subsequent analysis we used genes with appreciable PC_CT_ (|*Lp – Lo*|> 0.2 *Lo*) and EC_T_ (|*La – Lp*|> 0.2 *Lo*) (Ho & Zhang, 2018) and calculated the proportion of plastic genes with reinforced (TC > PC_CT_) or reversed (TC < PC_CT_) changes.

**Figure 2:**
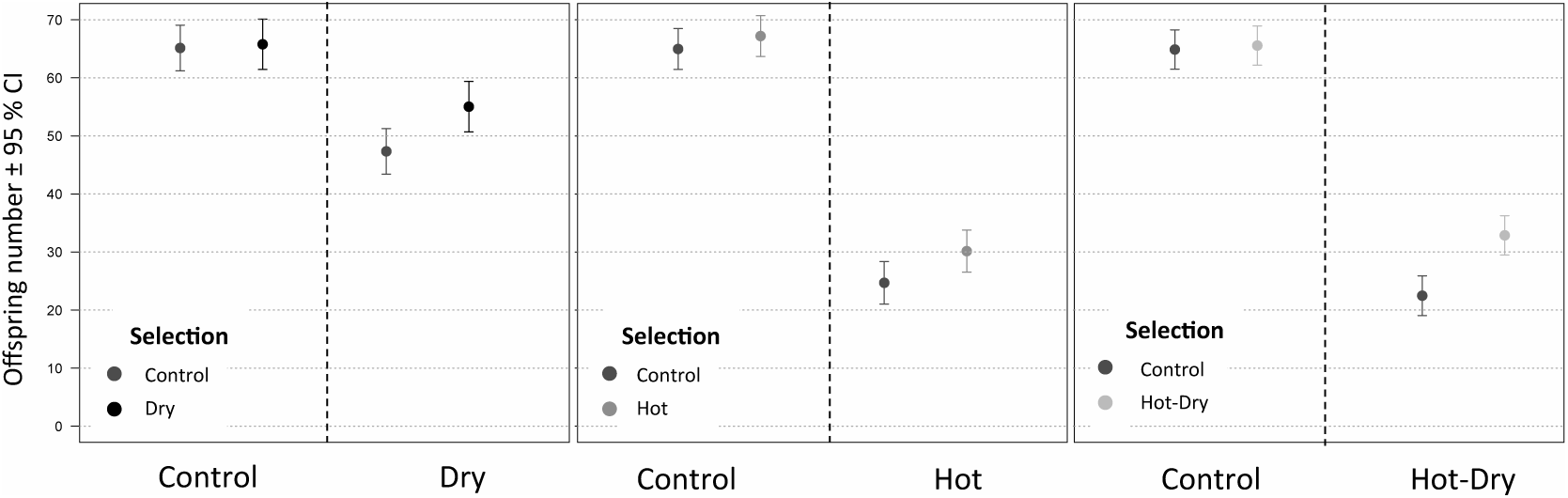
Offspring number of Control and selection lines under different conditions. Selection lines could adapt for 20 generations.

An alternative classification of plastic responses is to assess whether PC_CT_ brings expression levels closer to the new optimum (*La).* In this case, the total change TC would be larger than EC_T_ (Figure 2C). Such a pattern can occur even if PC_CT_ and EC_T_ are in opposite directions and classified as reversion (Figure 2C), but may indicate that ancestral plasticity was beneficial. A PC_CT_ that moves *Lp* further away from *La* is strongly indicative of maladaptive plasticity.

To confirm that our results were not sensitive to the applied cutoff (20% *Lo*), we repeated the analysis with a cutoff of 50% *Lo* and without any cutoff (Supporting information, Figure S3.1 A-H). Recently, it was pointed out (Mallard et al. 2018; Ho and Zhang 2019) that an excess of reversions relative to reinforcements is expected to be observed due to a statistical artefact that cannot be completely removed by permutation tests. We therefore applied an additional parametric bootstrapping test as proposed by Ho and Zhang (Ho and Zhang 2019), to compare proportion of reversions and reinforcements (Supporting information, Figure S3.1 J-L).

To better understand the relationship between the within-line proportions of reversed genes and proportions of genes with *La* closer to *Lo* with adaptation, we calculated the Spearman correlation between the proportions of reversed expression changes (or *La* closer to *Lo* respectively) and mean offspring number in seven selection lines in HD. We focused on HD because it was the most extreme environment with the strongest decline in offspring number. To test for significance, we used permutations: Mean offspring numbers were randomly assigned to lines and correlation was calculated again. Proportion of permutations with a correlation coefficient exceeding the observed value gave the respective *P*-value.

## Results

### Fitness assay showed evolutionary adaptation

We found that selection lines had a higher offspring number in their native condition compared to CT-lines (Dry: F_1,14_ = 9.20, *p* = 0.009; Hot: F_1,16_ = 4.78, *p* = 0.044; Hot-Dry: F_1,16_ = 23.51, *p* = 1.786E-04), confirming that adaptation had occurred (Figure 2). In contrast to treatment conditions, there was no difference in offspring number between CT- and selection lines under CT conditions (CT-lines: 64.89 [61.49, 68.29]; D-lines: 65.76 [61.26, 70.25], H-lines: 67.41 [63.76, 71.06], HD-lines: 65.56 [62.17, 68.94]) (Figure 2). Using three additional mixed models, we compared how lines from different selection regimes responded to treatments. We found significant negative effects for all stress treatments (D: F_1,23_ = 45.37, *p* = 6.85E-07; H: F_1,28_ = 507.68, *p* < 2.200E-16; HD: F_1,28_ = 553.06, *p* < 2.200E-16) (Figure 2). Interaction between selection and treatments, i.e. whether the response to the treatment was different depending on selection regime, was significant for HD-lines (F_1,28_ = 9.39, *p*= 4.754E-03) and for D (F_1,23_ = 4.32, *p* = 0.049), but not for H (F_1,28_ = 0.57, *p* = 0.455). Interaction between selection regime, treatment and line type (normal/mixed) was not significant in any treatment indicating that mixing lines did not have an effect on adaptation. ANOVA tables as well as results of the linear mixed models are in supporting information S4.

### Evolution of gene expression

We found many more genes with expression divergence relative to Control (TC) than genes with evolved changes in the treatments (EC_T_) or in control conditions (EC_CT_) (Table 2). Increased temperature caused expression divergence in a larger number of genes than reduced humidity (Table 2). Most DE genes with significant TC also had significant ancestral plasticity PC_CT_ (Dry: 33.9%; Hot: 72.9%, Hot-Dry: 82.2%), but only a minority showed significant evolutionary change EC_T_ (Dry: 4.9%; Hot: 1.1%; Hot-Dry: 1.0%, see also Supporting information Figure S3.2). About half of the DE genes with EC_T_ in Hot and Hot-Dry then did not diverge significantly from their ancestral expression levels in Control (TC ∼ 0). Overall, TC had a larger log2-fold change contribution from plastic changes (PC_CT_) than evolved changes in Hot and Hot-Dry (median proportions PC_CT_/TC; Hot: 71.47%; Hot-Dry: 86.30%), but not in Dry (42.90%). Many genes had PC_CT_ > TC, especially in HD (Figure 3A). In these cases, EC_T_ was negative and reduced TC.

**Figure 3:**
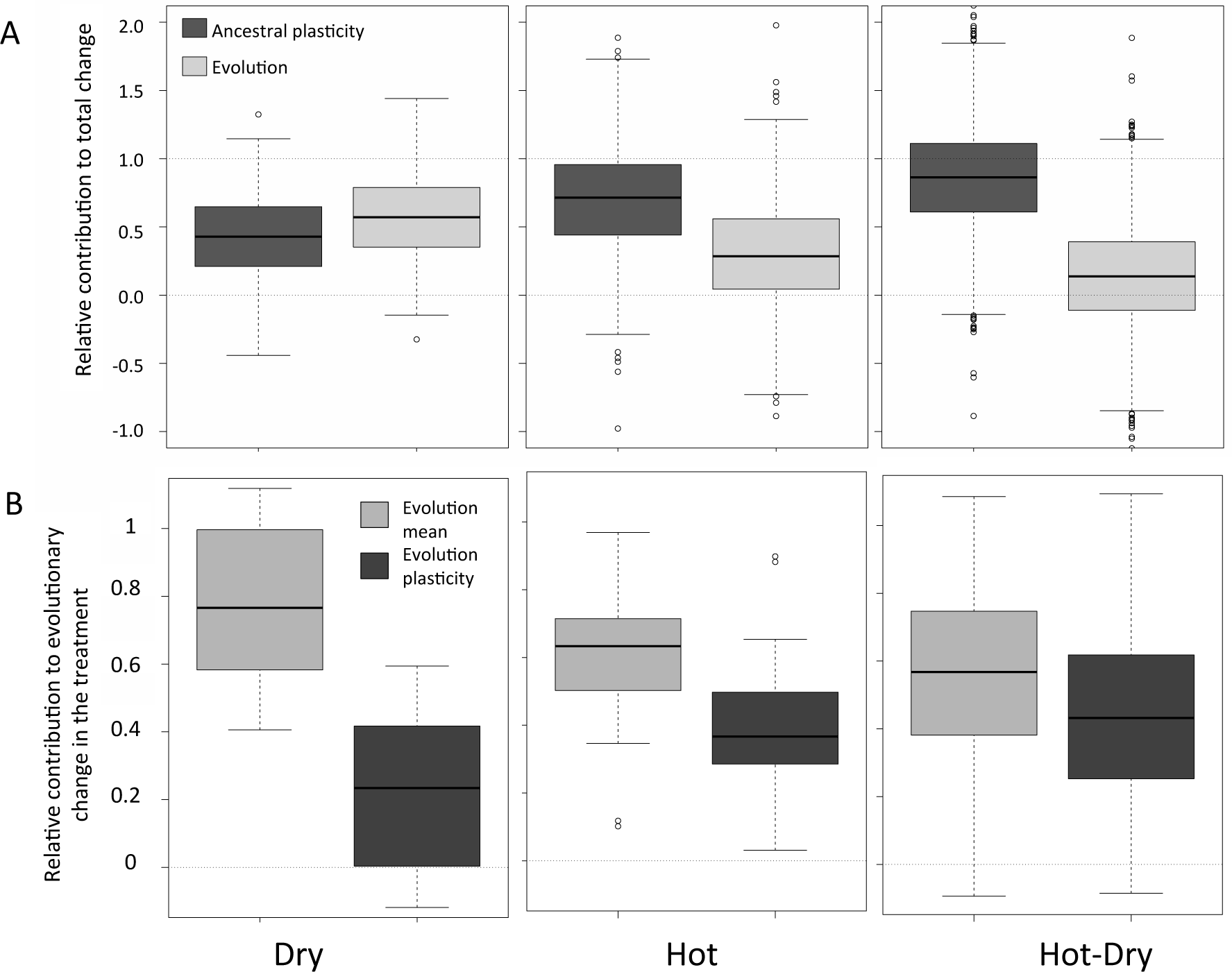
Relative contribution of ancestral plasticity and evolutionary change to total divergence for all genes showing significant total divergence in the DE analysis. Number of genes: Dry: 283, Hot: 1192, Hot-Dry: 2045 **(A)**. Relative contribution of changes in the mean (shift in the intercept of reaction norm) and changes in plasticity (different slopes of reaction norms) to evolutionary differences between control-lines and selection lines in the treatment. Only genes with significant differences in the DE analysis were included. Number of genes: Dry: 18, Hot: 25, Hot-Dry: 55 **(B)**.

**Table 2:** Significantly differently expressed genes. For evolved differences, control-lines and selection-lines are compared within condition (e.g. Dry-lines vs Control-lines in Dry). Plastic response gives the number of genes that changed expression between control and treatment conditions (e.g. Dry-lines in Dry vs Dry-lines in control). Different plasticity gives the number of genes with significant different plastic responses in lines from different selection regimes. Analysis was conducted using the R package limma (Ritchie et al., 2015).

|  |  | Dry | Hot | Hot-Dry |
| --- | --- | --- | --- | --- |
| Evolved difference in Control | down | 3 | 0 | 6 |
|  | up | 1 | 9 | 4 |
|  | **total** | **4** | **9** | **10** |
| Evolved difference in treatment | down | 9 | 8 | 22 |
|  | up | 9 | 17 | 33 |
|  | **total** | **18** | **25** | **55** |
| Plastic response Control-Lines | down | 209 | 1765 | 2451 |
|  | up | 156 | 1714 | 2200 |
|  | **total** | **365** | **3479** | **4651** |
| Plastic response Selection-Lines | down | 28 | 1417 | 1649 |
|  | up | 21 | 1381 | 1470 |
|  | **total** | **49** | **2798** | **3119** |
| Total change (Selection-lines in treatment vs Control-lines in Control) | down | 154 | 564 | 1016 |
|  | up | 129 | 628 | 1029 |
|  | **total** | **283** | **1192** | **2045** |
| different plasticity | **total** | **0** | **1** | **4** |

### Evolution of reaction norms

We were then interested to examine whether EC_T_ were due to a change in mean expression (shift in the intercept of reaction norms) or to a change in plasticity (different slopes of reaction norms, see Figure 3). First, we found a significant correlation between EC_CT_ and EC_T_, in all three treatments (permutation tests: *P* < 0.0001) (Table 3, Supporting information Figure S2). This indicates that for a majority of genes, evolution in the past 20 generations affected their mean expression levels by shifting the intercept of reaction norms (see Figure 1D). Second, we also found a highly positive correlation between PC_CT_ and PC_Sel_ (Figure Supporting information S2A), indicating that plasticity was mainly preserved during adaptation to new conditions. A permutation test showed that this correlation was not significantly different from correlations obtained from permuted datasets (Table 3), where samples were randomly assigned to selection regimes, suggesting that the treatments did not have specific effects on plasticity. When we quantified the relative contributions of changes in the mean versus changes in plasticity to EC_T_, we found that evolution of the intercept explained more evolutionary divergence than evolution of the slope of reaction norms, especially in Dry (Figure 3B).

**Table 3.** : P-values obtained from permutation tests (10,000 permutations). Samples were randomly assigned to either control or treatment selection and differential expression analyses were repeated. Significance was assessed by calculating the proportion of permutations with more extreme values than the observed one. Control (CT) conditions: 33°C, 70% relative humidity r.h. Conditions in treatments: Dry (D): 33°C, 30% r.h.; Hot (H): 37°C, 70% r.h.; Hot-Dry (HD): 37°C, 30% r.h.

|  | Selection |  |  |
| --- | --- | --- | --- |
|  | D | H | HD |
| CT-Lines have more genes with significant plastic responses compared to adapted selection lines | 0.2981 | 0.3037 | **0.0325** |
| Magnitude of plastic response is higher in CT-Lines | **0.0493** | 0.1676 | **0.0491** |
| Individuals from different selection regimes show lower similarity in plastic responses<sup>†</sup> | 0.8608 | 0.3914 | 0.5217 |
| Number of genes with significant differences in expression levels is higher in treatment than in CT conditions | **0.0095** | **0.0003** | **0.0031** |
| Differences in expression levels in CT conditions and treatment are correlated | **< 0.0001** | **<0.0001** | **< 0.0001** |

†: observed correlation of plastic responses of CT- and selection lines would be lower compared to results of permutations

### Evolution of plasticity in DE genes

We found only five genes with significantly evolved plasticity in the DE analysis, after correcting for multiple testing. Although plasticity seemed to be preserved overall, we found evidence that it evolved among DE genes. The DE analysis revealed a reduced number of plastic genes in selection lines compared to plastic genes in CT-lines (Table 2), although the difference was only significant in HD (Table 3). It further showed that ancestral plasticity PC_CT_ was larger than evolved plasticity PC_Sel_, in D and HD (Table 3), indicating a reduction of the magnitude of plastic responses after adaptation. We found the same outcome when focusing on genes with significant EC_T_, in HD (Mann-Whitney U test: U = 2133, *P* = 2.102E-04; median: CT-lines: 0.25, HD-lines: 0.13), but not in D (U = 144, *P* = 0.584; median: CT-lines: 0.10, D-lines: 0.14) or H (U = 386, *P* = 0.158; median: CT-lines: 0.21, H-lines: 0.12) (Figure 4).

**Figure 4:**
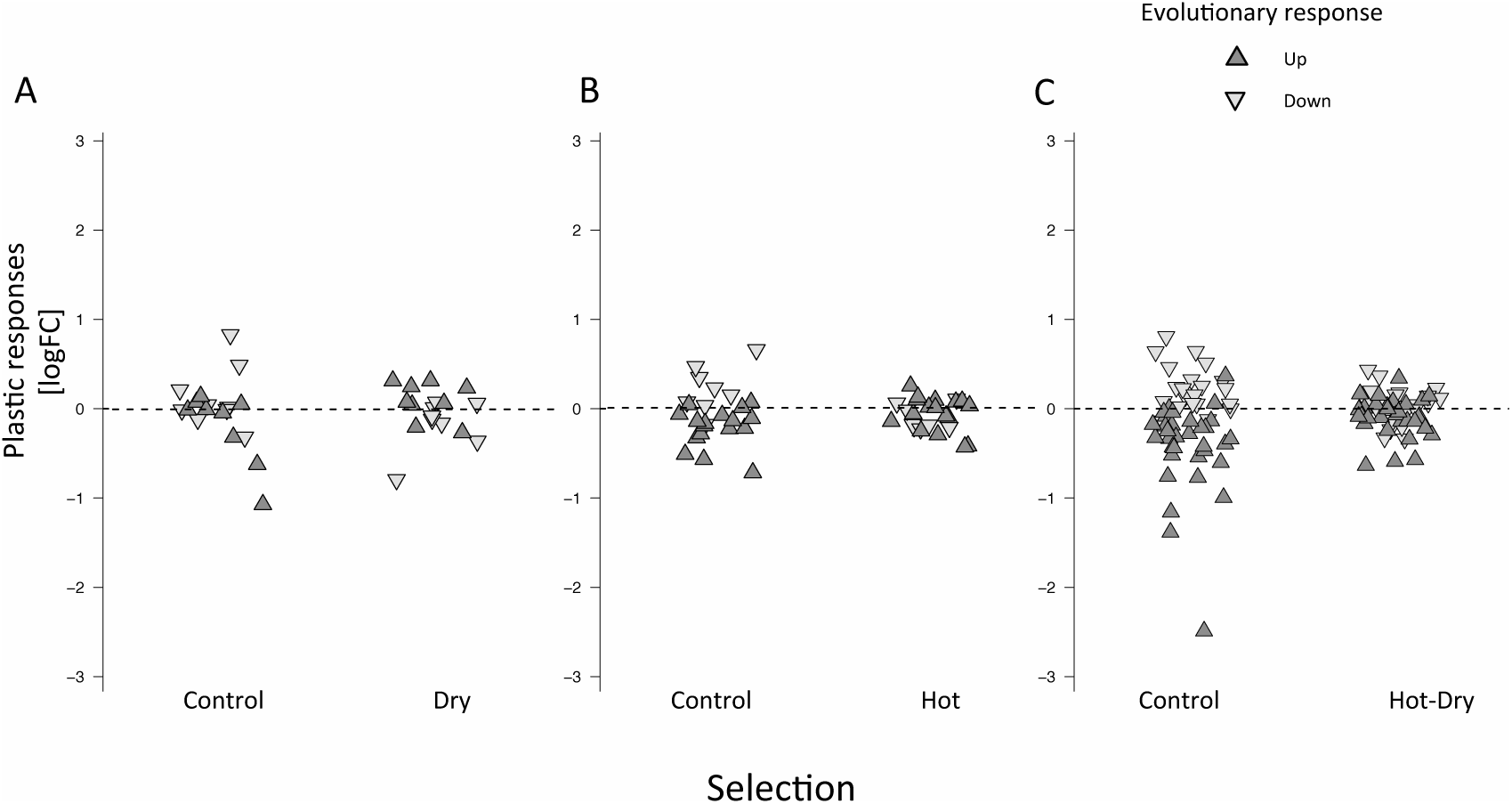
Plastic responses in control-lines and adapted selection-lines of genes showing significant evolutionary changes in expression in the treatments Dry **(A)**, Hot **(B)** and Hot-Dry **(C)**. Plastic responses in adapted selection lines are weaker (smaller logFC). Furthermore, plastic changes are mainly opposite to evolved changes. Genes that evolved to lower expression in the treatments (represented in lightgrey) show positive plastic changes in control-lines, i.e. they are up-regulated. Genes that evolved to higher expression levels (in darkgrey) are down-regulated in the non-adapted Control-Lines.

### Effect of plasticity on evolutionary responses

We could not find evidence that plasticity in general impeded evolution. In the DE analysis genes with a significant ancestral plastic response were overrepresented among the evolved DE genes in H and HD (H: χ2 = 4.51, df = 1, *P* = 3.38E-02; HD: χ2 = 9.71, df = 1, *P* = 1.84E-03), but not in D (*P* = 0.113 Fisher’s exact test). Furthermore, when we analysed each selection line separately and compared their expression levels to CT-lines, the proportion of genes exhibiting substantial PC_CT_ (|*Lp* – *Lo*| > 0.2 *Lo*) were also overrepresented among evolved genes with EC_T_ (|*La* - *Lp*| > 0.2 *Lo*) (Table S5.1).

Maladaptive plastic changes may be reverted by evolutionary changes when the plastic response overshoots the optimum expression level in a new environment. Among the genes with significant EC_T_ in the DE analysis, we found that PC_CT_ was generally in opposite direction (Figure 4). Enrichment tests revealed that genes with positive EC_T_ in the DE analysis were overrepresented among genes with negative PC_CT_ in H and HD (Fisher’s exact test D: *P* = 0.076; H: *P* = 0.002; HD: *P* = 9.26e-06), and genes with negative EC_T_ were overrepresented among genes with positive PC_CT_, in H and in HD as well (Fisher’s exact test D: *P* = 0.073; H: *P* = 0.021; HD: *P* = 0.049)(Figure 4). In almost all genes that showed significant EC_T_ and PC_CT_, the responses were in opposite direction (Dry: 2 out of 2 genes; Hot: 12 out of 13; Hot-Dry: 33 out of 34). We also found negative correlations between PC_CT_ and EC_T_ in all treatments, although significant only in H and HD, but not in D (Supporting information, Figure S3.3).

When we considered mean expression levels per line with PC_CT_ > 20% *Lo* and EC_T_ > 20% *Lo*, we obtained similar results as in the DE analysis. Reinforcements were present but less frequent than reversions in all treatments (proportions of plastic responses with reversion/reinforcement: Dry: 48.5 ± 1.0% / 34.3 ± 0.8 %, *P* < 1.151e-07 (binomial test); Hot: 44.6 ± 0.9% / 28.8 ± 1.2%, *P* = 4.369E-13; Hot-Dry: 43.6 / 27.1 *P* < 2.2e-16; Figure 5A). The reversed genes represented 34.1 ± 0.8% (Dry), 40.0 ± 0.9% (Hot), and 42.96 ± 2.1% (Hot-Dry) of the evolved genes. In contrast, 24.1 ± 0.6% (Dry), 25.8 ± 1.1% (Hot), 26.80 ± 1.6% (Hot-Dry) of evolved genes showed reinforcement.

**Figure 5:**
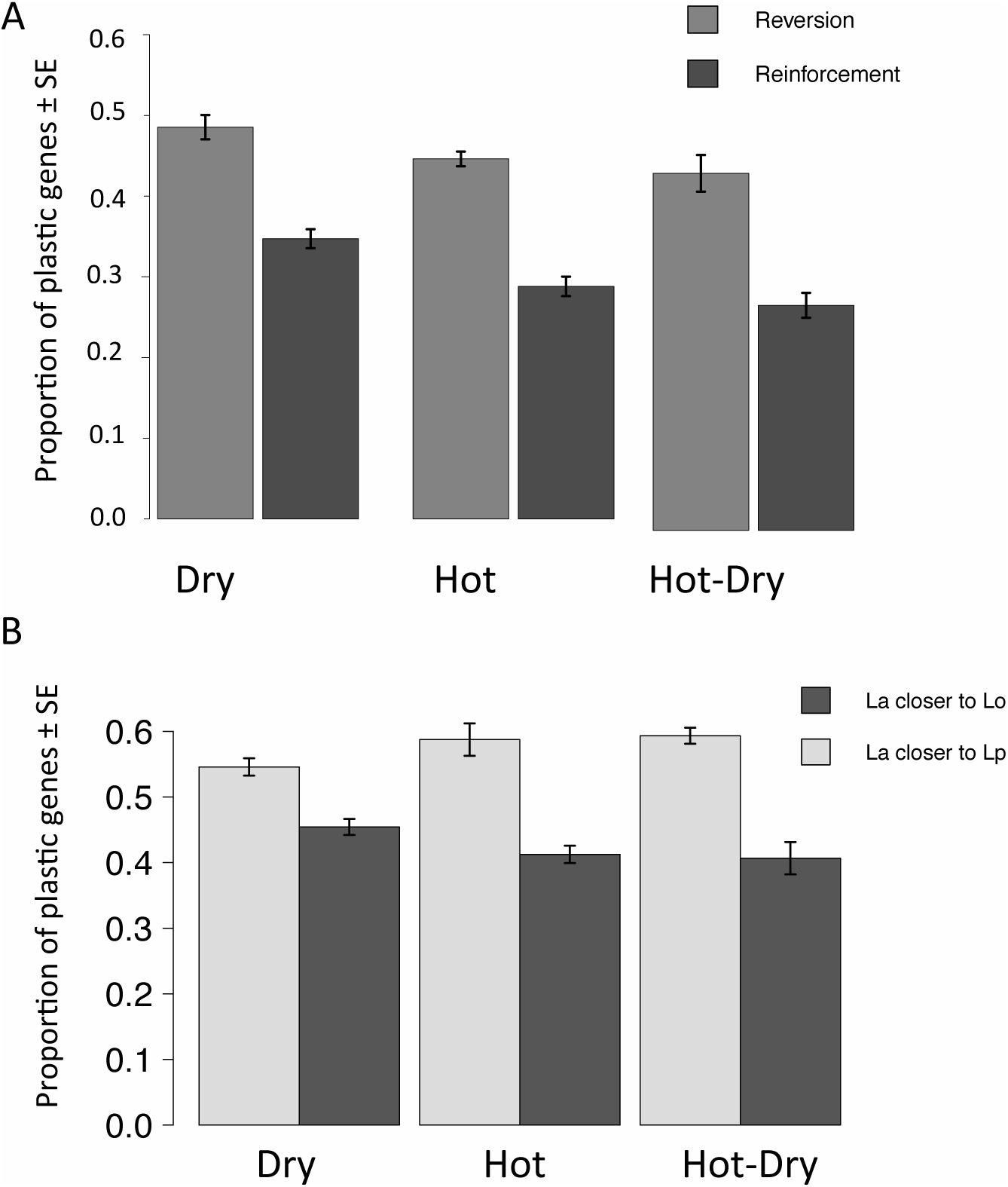
Comparison of plastic and evolutionary changes in gene expression in response to three environmental conditions. **A:** Proportion of genes (average over selection lines) showing a reversion or reinforcement of ancestral plasticity during evolution. **B:** Proportion of genes with expression levels after evolution (adapted stage La) closer to original levels (Lo, i.e. Control-lines in control conditions), or closer to plastic levels (Lp, i.e. expression levels of Control-lines in treatment). Only genes exhibiting substantial plastic changes (|Lp-Lo| > 20% of Lo) as well as evolutionary changes (La-Lp| > 20% of Lo) were used for analysis.

Different cut-off values did not influence these outcomes (Figure S3.1). Parametric bootstrapping as an additional test (Ho & Zhang, 2019), confirmed our results (Supporting information, Figure S3.1). We also found that a large proportion of ancestrally plastic genes had *La* closer to *Lo* than to *Lp* (Dry: 45.4 ± 1.1%; Hot: 41.2 ± 1.2%; Hot-Dry: 40.7 ± 2.5 %)(Figure 5B). Finally, substantial proportions of genes with EC_T_ did not show PC_CT_ (Dry: 41.8 ± 0.7%, Hot: 34.6 ± 0.8%, Hot-Dry: 30.3 ± 1.0 %).

In case of genetic assimilation, i.e. loss of plasticity in selection line and continuous expression of the induced phenotype, evolutionary differences would not be visible in treatment condition. We calculated the mean expression levels of selection lines in Control and defined genes as showing assimilation if they met three criteria: no PC_Sel_ (absolute difference between selection lines in CT and in treatment < 20% *Lo*), PC_CT_ > 20% *Lo*, and *Lp* ∼*La* (absolute difference < 20% *La*). The proportion of PC_CT_ genes showing genetic assimilation was small in all conditions (Dry: 6.8 %, Hot: 10.3 %, Hot-Dry: 10.3 %; Supporting information Table S5.2).

### Proportions of reversed plastic responses and association with fitness

To gain a better understanding of the adaptive value of the changes of expression levels in the evolved lines, we tested for an association between within-line proportion of reversed or reinforced plastic responses and the average fitness of the lines in the HD treatment. We found that lines with a higher proportion of reversions had a higher average offspring number (correlation: 0.82, *P* = 0.012, Figure 6A) and we found a negative but non-significant correlation between fitness and reinforcements (correlation: −0.43, *P* = 0.85). When we tested for an association between fitness and proportion of ancestrally plastic genes with *La* closer to *Lo*, we also found a positive correlation (correlation: 0.86, *P* = 0.006, Figure 6B). Overall, better adapted lines (higher fitness in HD) showed a higher proportion of reversed ancestral plasticity and these plastic genes were more similar to the original expression levels of CT-lines in CT. Performing the analysis with gene expression data in H and D provided similar correlations, although not significant because of lower sample sizes (Supporting information, Table S5.3).

**Figure 6:**
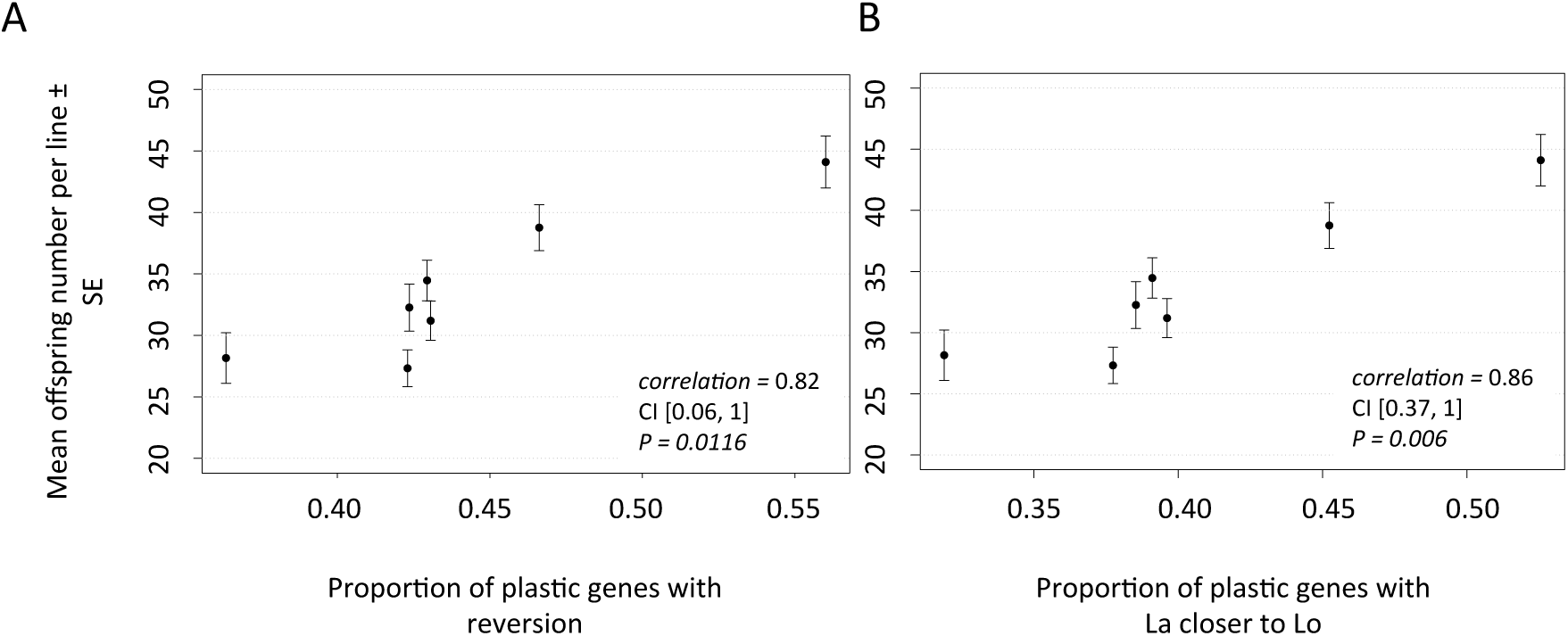
Relationship between plastic and evolved changes in gene expression in response to hot-dry conditions. Expression levels of Control-lines in control conditions represent the ancestral stage Lo, Control-Lines in hot-dry conditions the plastic stage Lp and adapted Hot-Dry-lines in hot-dry conditions the adapted stage La. **A:** Relationship between the proportion of reversed plastic responses and mean fitness (=offspring number) per Hot-Dry line. **B:** Relationship between proportion of genes with La closer to Lo and mean fitness. Only genes exhibiting substantial plastic changes (|Lp-Lo| > 20% of Lo) as well as evolutionary changes (La-Lp| > 20% of Lo) were used for analysis. P-values of the spearman correlations were obtained by 10,000 permutations. 95 % Confidence intervals are based on a non-parametric bootstrap test.

## Discussion

We studied plastic and evolved responses in gene expression of *T. castaneum* in response to three new environmental conditions (Dry, Hot, Hot-Dry). After 20 generations of experimental evolution, we were able to detect adaptation and found significant evolutionary changes in expression levels. Comparing evolutionary changes with ancestral plastic responses showed that a reversion of plasticity was most frequent (> 95 % genes with significant plastic changes in DE analysis; > 40 % of genes with substantial plastic changes, i.e. changes higher than 20% of ancestral levels). The number of genes where ancestral plasticity was reinforced by evolution was significantly smaller (DE analysis: <5%, 27-34 % of genes with substantial plastic changes) and a smaller proportion of genes (7-10 % of substantially plastic genes) showed expression patterns consistent with genetic assimilation. A high proportion of the originally plastic genes evolved to expression levels that were closer to control levels than to ancestrally plastic levels. Although the proportion of non-reversed plastic genes was still high, positive associations between fitness and proportion of reversions, and compensated plasticity (expression levels closer to control levels) respectively, suggest that ancestral plasticity was maladaptive for a majority of responding genes. Although plasticity showed a high degree of preservation in terms of number of responding genes and direction of the response, we found evidence that selection lines evolved a reduced plasticity and thus partly compensated the maladaptive ancestral response. We were further able to show a positive association between the proportion of reversed plastic responses and adaptation (mean fitness per line) in the most stressful treatment Hot-Dry.

### Maladaptive plasticity

Different patterns describing the relationship between plastic and evolved changes in gene expression have been documented. It was suggested that plasticity might help populations to persist after environmental change, or to colonize new habitats by bringing phenotypes closer to the new optimum. Studies found support for this hypothesis by showing that plastic responses of non-adapted individuals diminished differences to native populations (Lohman, Stutz, & Bolnick, 2017; Mäkinen, Papakostas, Vøllestad, Leder, & Primmer, 2016). Adaptive plasticity can also be indicated when plastic and evolutionary responses are in the same direction (Li, Li, Song, Wang, & Zhang, 2017), or when plasticity is higher in adapted populations (Hasan et al., 2017; McCairns & Bernatchez, 2009), suggesting that most plastic individuals were favored by selection. However, there are also examples for the reversed pattern suggesting that plasticity was maladaptive. In wild populations of *Fundulus heteroclitus*, evolved changes to different temperatures were opposite to plastic responses of the ancestral population (Dayan et al., 2015). *Rhagoletis* flies shifting to a new host fruits showed evolutionary responses opposite to plasticity of non-adapted species (Ragland et al., 2015). Experimental evolution studies found countergradient evolution in *Drosophila* adapting to different diets (Huang & Agrawal, 2016; Yampolsky et al., 2012) and in guppies adapting to low predation environments (Ghalambor et al., 2015). A comparative study (Ho & Zhang, 2018) analyzing data of multiple experimental evolution suggested that reversions of gene expression changes might be a general pattern during adaptation.

Our study fits with these previous observations. We found a higher proportion of reversions than reinforcements in all conditions indicating mostly maladaptive plasticity. An alternative explanation for the prevalence of reversions without maladaptive plastic responses would be that CT-lines exhibited a response in the right direction, but overshot an optimum expression level (Figure 1C). Fine-tuning during long-term adaptation could then lead to a partial reversion of the plastic response. We took this possibility into account by not only focusing on a reversion of plasticity, but also testing whether plastic changes brought expression levels closer to the adapted stage *La* than to original stage *Lo.* If, in contrast, we see that *La* is closer to the original level (CT lines in CT conditions), it indicates that plastic responses were maladaptive since they moved expression levels further away from the new optimum and became compensated during evolution (see Figure 1C). We found that adapted lines showed a high proportion of expression levels closer to their ancestral level *Lo* than to *Lp*. We could further show that this proportion is positively associated with higher fitness per selection line. We also found strong positive associations of within-line proportions of reversions with mean reproductive output, indicating a possible fitness advantage to reversions.

There was still a large proportion of plastic genes that did not show reversion. They are either close to the levels of adapted lines, or showed reinforcement. The first case might indicate that plasticity prevented evolution by matching the new optimum. Reinforced plastic changes could be examples of adaptive plasticity. However, correlation between proportion of reinforcements and fitness was not significant and negative. The observed positive correlation between proportion of reversed plastic genes and mean fitness per line in HD rather suggests that reversions were favoured during adaptations. Reversions might become more pronounced after more generations, once evolution had sufficient time to further reverse maladaptive ancestral plastic responses. Our selection lines still show a strong reduction in offspring number compared to control levels suggesting further potential to adapt.

### Evolution of reaction norms

The ancestral maladaptive plasticity can be compensated by shifts in the intercept or changes in the slope of reaction norms. Both are not mutually exclusive and can occur together in the same trait. We were aiming to quantify their relative importance for evolutionary responses in transcriptomes.

Gene expression studies so far provided mixed results regarding the evolution of plasticity. *Drosophila* populations adapted to different temperatures showed local adaptation, but there was no evidence for evolution of thermal reaction norms of different transcripts and changes affected mainly expression mean (Clemson, Sgrò, & Telonis-Scott, 2016). Experimental evolution studying *Drosophila* under variable diets found no significant changes in plasticity (Yampolsky et al., 2012) or less than expected (Huang & Agrawal, 2016). In contrast, other studies found differences in temperature responses between tropical and temperate *Drosophila* populations (Levine, Eckert, & Begun, 2011; von Heckel et al., 2016). Other examples for differences in genes expression plasticity between adapted and non-adapted populations include temperature (Morris et al., 2014) and salinity (Gibbons, Metzger, Healy, & Schulte, 2017; McCairns & Bernatchez, 2009) responses of marine and freshwater sticklebacks, temperature response of killifish populations from different latitudes, as well as plastic responses to toxic hydrogen sulphide (H_2_S) of fish population from H_2_S rich springs versus non-toxic springs (Passow et al., 2017). There was no consistent pattern regarding the direction in which plasticity evolves: In some cases adapted population showed an increase in plasticity (Morris et al., 2014), in other cases plasticity was reduced (Huang & Agrawal, 2016; Ragland et al., 2015; von Heckel et al., 2016) or reduction and enhancement of plasticity were equally frequent (Gibbons et al., 2017; Yampolsky et al., 2014). Overall, there is evidence in multiple species that expression plasticity of some genes can evolve. However, even in some of these studies reporting evolved plasticity (Dayan et al., 2015; Gibbons et al., 2017; Morris et al., 2014) the number of transcripts with significant changes in the mean was much higher than transcripts with changed plasticity and large parts of the plastic responses showed a high degree of preservation.

In accordance with these previous findings we found that changes in the mean contributed more to the observed expression differences in the treatments than changes in plasticity. A possible reason might be that genetic variation in mean expression was higher than genetic variation in plasticity. In addition, we did not select for changes in plasticity directly since the conditions in the treatments were constant. Selection was therefore on expression levels in the treatment and only indirectly on plasticity. Plasticity could evolve if mean expression levels were genetically correlated with plasticity. Although we did not directly test for such correlations, we found evidence for evolution of plasticity of the DE genes, that is of the genes with significant changes in expression level between CT and treatment conditions. Although plastic responses showed a high degree of preservation in terms of affected genes and direction, we found evidence for evolutionary changes in the magnitude of plastic responses, i.e. the slope of the reaction norm.

### Why can plasticity be maladaptive?

New stressors might disturb homeostasis resulting in inappropriate responses and long-term adaptation therefore restores ancestral phenotypes by genetic changes, referred to as genetic compensation (Grether, 2005) or counter-gradient variation (Conover et al., 2009). However, in our study we applied relatively mild stressor treatments, i.e. individuals were able to survive and reproduce. Drought and heat are also stressors, which *T. castaneum* had experienced in the past (Sokoloff, 1972), so there had probably been selection on plastic responses to be beneficial. However, plastic responses might be optimized for a short-term exposure: Allocation of resources from reproduction to protection might increase survival probability and allow individuals to continue reproduction as soon as the stress has disappeared, but this response becomes maladaptive during continuous exposure and should therefore be under negative selection. Expression of stress related genes is in general accompanied by a down-regulation of genes involved in growth and reproduction due to an allocation of resources (Schwenke, Lazzaro, & Wolfner, 2016; Sokolova, 2013). A well-studied example are heat shock proteins (hsp). Hsp are well known for their protective function and to be crucial for survival (Feder & Hoffman, 1999), but it was also shown that their expression comes at a cost (Feder et al., 1998; Sørensen, Kristensen, & Loeschcke, 2003). Accordingly, it was often found that hsp expression in populations adapted to warmer climates is lower compared to non-adapted populations (Fangue, Hofmeister, & Schulte, 2006; Narum & Campbell, 2015; J G Sørensen, Dahlgaard, & Loeschcke, 2001). In general, other protection mechanisms independent of ancestral plasticity may arise during long-term adaptation (e.g. enzymes, which are more stable at high temperature) and make the costly stress response expendable.

An alternative explanation for the reduced plasticity in adapted lines is that the signal responsible for eliciting the plastic responses is based on any kind of damage (e.g. deformations in macromolecules, membrane lipids, proteins, and DNA) caused by heat or stress in general (Kültz, 2005). Higher resistance in adapted lines might shift the inducing thresholds, i.e. the temperature when damages occur and stress response is induced (Sikkink, Reynolds, Ituarte, Cresko, & Phillips, 2014) above the levels we applied in the treatments.

Interestingly, we found no differences in fitness between lines from different selection regimes under CT conditions (Figure 2). We could detect some genes with different expression levels in the treatments and found in general a correlated change in expression levels in control conditions, but this did not affect offspring number. It indicates a lack of fitness trade-offs, where alleles providing a fitness advantage in one environment (treatment) are detrimental in another (CT). Together with the observation that selection lines evolved to bring expression closer to ancestral CT-levels, it suggests that for many genes the optimal expression level is not different between conditions. They might be involved in processes important for maintenance and reproduction. Under stress, limited resources have to be invested into protection, that are then not available for reproduction (Sokolova, 2013). Long-term adaptation should then work to restore control levels that are likely to be optimized for highest reproductive output and to reduce costly stress responses resulting in improved canalization of traits associated with fitness (Stearns & Kawecki, 2006). Canalization, i.e. robustness against environmental variation, was found previously in gene expression adaptation (Levine et al., 2011; Shaw et al., 2014; von Heckel et al., 2016). Genetic differences between control and selection lines that are responsible for adaptation to the treatments did not have an effect in control conditions and thus represent cryptic genetic variation (Gibson & Dworkin, 2004). They might either concern genes that are not expressed in control conditions, or represent changes neutral under control conditions.

### Potential caveats

The number of genes with significant plastic changes in the DE analysis was much higher compared to genes showing evolutionary changes. One possible explanation would be that adaptive plasticity prevented evolution. If the plastic responses matched the optimum, no genetic changes in the selection lines are expected to occur. However, when we analysed each line separately and considered a gene as evolved if the mean difference between *La* and *Lp* was more than 20% of the *Lo*, we found approximately the same number of genes with EC_T_ and PC_CT_ (Supporting information Table S5.1).

In the DE analysis in *limma* we did not analyse each line separately but treated them as biological replicates. Since lines were split across conditions comparisons between conditions, i.e. plastic changes, can be made within lines. They should thus be more precise and statistical power should be higher than comparisons between selection regimes, i.e. evolved changes, that have to be made between lines. Differences between lines from the same selection regime lower the ability to obtain significant evolutionary changes. These differences can arise from genetic drift. Since our population size was relatively small (120 individuals per line) this might have been an important factor. Another explanation is that lines from the same selection regime differed how exactly they improved their fitness in the respective treatment. Since fitness is a highly polygenic trait, the genes contributing to a fitness increase may not be the same in different lines (see Barghi et al., 2019). For the most extreme treatment HD, where we sequenced seven lines, we further found considerable differences in fitness between the lines, suggesting that not all of them were at the same stage of adaptation. It is therefore not surprising that expression levels did not evolve in the same way among lines.

The DE analysis in *limma* requires that a gene shows similar changes in all replicate lines, and is therefore more conservative. If the main interest of a study is to identify promising candidate genes for future more detailed analyses it is the appropriate approach to keep FDR as low as possible. In contrast, if the focus is more on general patterns, a less stringent analysis using mean expression levels can give us a more complete picture. Since genetic drift is random it cannot explain the observed excess of reversions over reinforcements.

Although we demonstrated that gene expression changed during evolution, it is not clear whether these changes are the cause of an increased fitness in these conditions or whether they are rather the consequence of adaptation and being less stressed. One disadvantage in studying whole transcriptomes is that not all responding genes might be of functional importance but are correlated to other adaptive changes. High intercorrelations within the transcriptome (Ayroles et al., 2009; McGraw et al., 2011) might lead to correlated responses in many other genes. Furthermore, observed evolutionary changes might be caused by indirect selection and other mechanisms, e.g. changes in protein structure of enzymes, were responsible for adaptation of selection lines. Future studies that manipulate expression and test for correlated changes in offspring are needed to confirm adaptive value of expression changes.

## Conclusions

We found that genes with the strongest plastic responses showed evolutionary changes in opposite direction suggesting that ancestral plasticity was maladaptive for long-term adaptation. In the most stressful treatment, selection lines with higher fitness show a higher proportion of reversions and a higher proportion of originally plastic genes that are closer to ancestral expression levels. Differences between adapted lines and CT-lines in the treatment were mainly due to a change in mean expression (i.e. shift in the intercept of reaction norms), while plasticity was preserved in terms of affected genes and direction of change. However, we found that a part of the differences in the treatments can be explained by a reduction in the magnitude of plasticity in adapted lines. Our results add to growing evidence that plasticity and evolution are often in opposite direction and maladaptive plastic responses might increase strength of selection. In contrast to previous studies, including fitness data allowed us to give evidence for adaptation and in the most stressful condition we were able to show an association between reversion of plasticity and adaptation. Similar results in all three stress treatments indicated that these findings may represent a general pattern of gene expression adaptation.

## Data Accessibility

RNA-seq reads will be available at GEO. Fitness data will be available at Dryad.

## Author Contribution

FG and ELK designed experiment. ELK conducted experiment, laboratory work and analysed the data. FG and ELK wrote the manuscript.

## Acknowledgements

This work was supported by the Swiss National Science Foundation, grants PP00P3_1144846 and PP00P3_176965 to FG. We thank Sonja Sbilordo, Sara Meier, Cilgia Lippuner, Tim Schoch and Tim Emmenegger for helping during the fitness assay, Elke Karaus, Martina Berchtold, Encarnación Lozano for RNA extraction and quality control, Lucy Poveda and Maria Domenica Moccia for advice in library preparation and sequencing, Lennart Opitz for helping to process the RNA-seq data.

