## Supporting information S3 for "Restoring ancestral phenotypes by reduction of plasticity is a general pattern in gene expression during adaptation to different stressors in *Tribolium castaneum*"

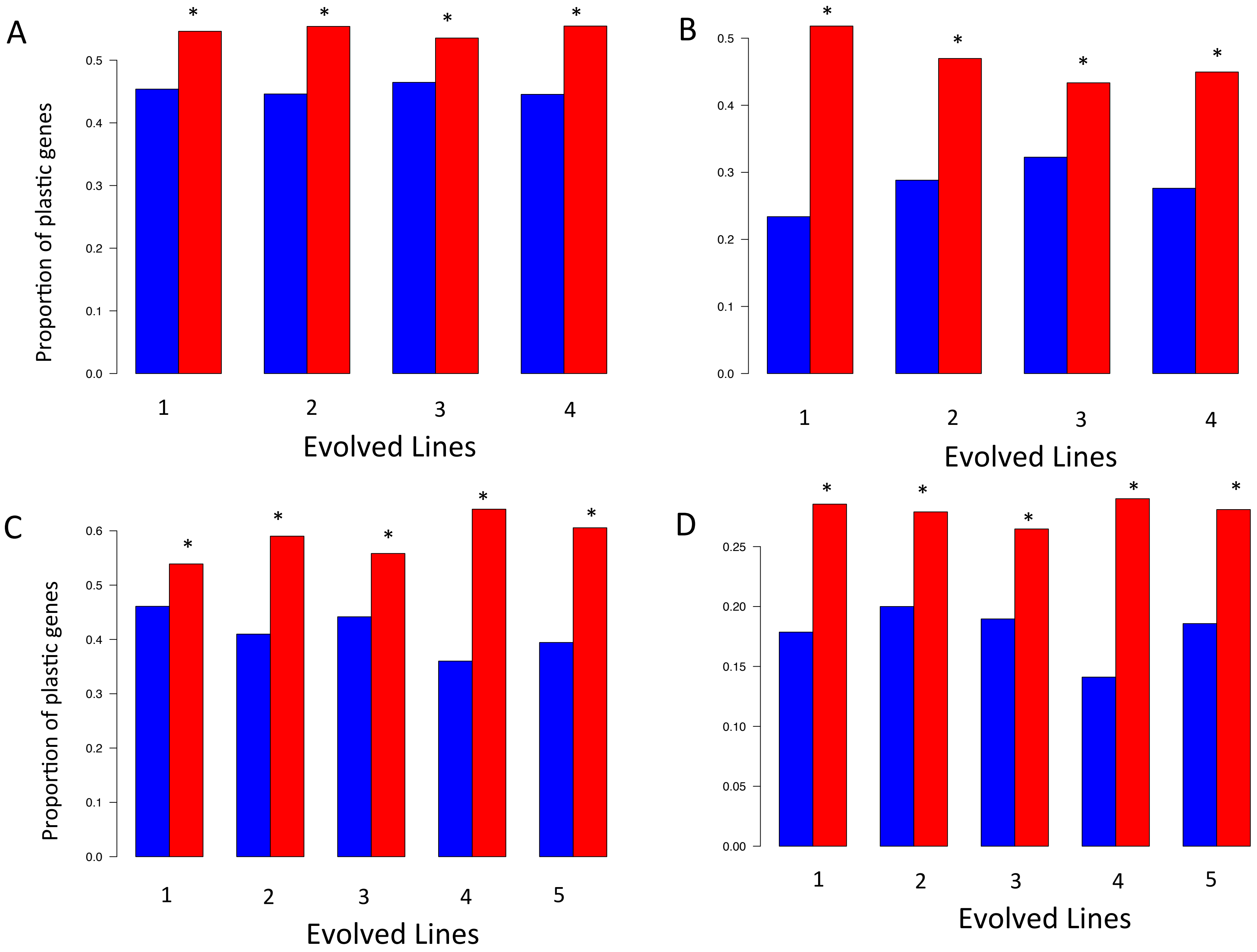
A
B
C
D
Proportion of plastic genes
Evolved Lines
0.0
0.1
0.2
0.3
0.4
0.5
0.6
0.00
0.05
0.10
0.15
0.20
0.25
1
2
3
4
5
*

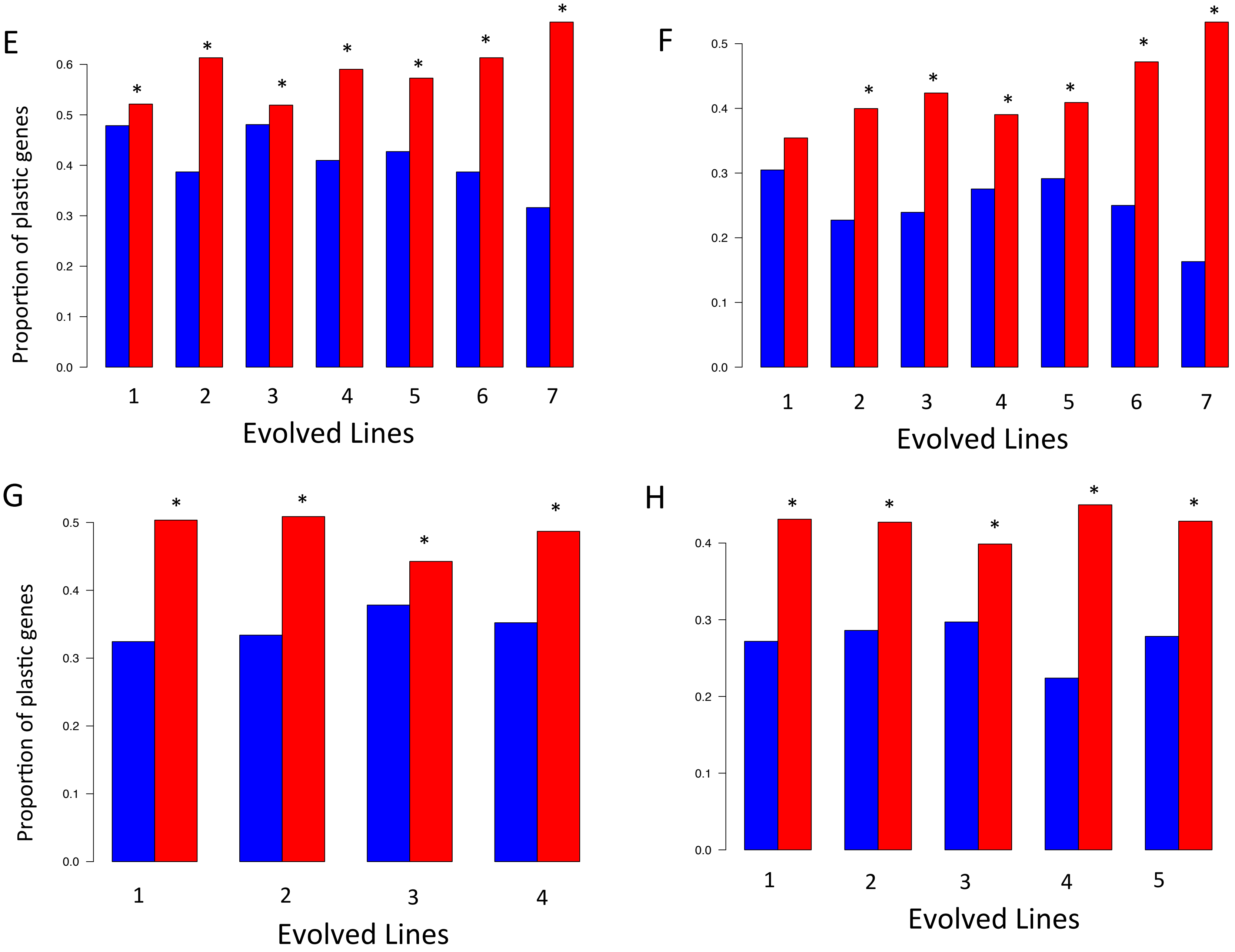
E
F
G
H
Proportion of plastic genes
Evolved Lines
0.0
0.1
0.2
0.3
0.4
0.5
0.6
1
2
3
4
5
6
7
*

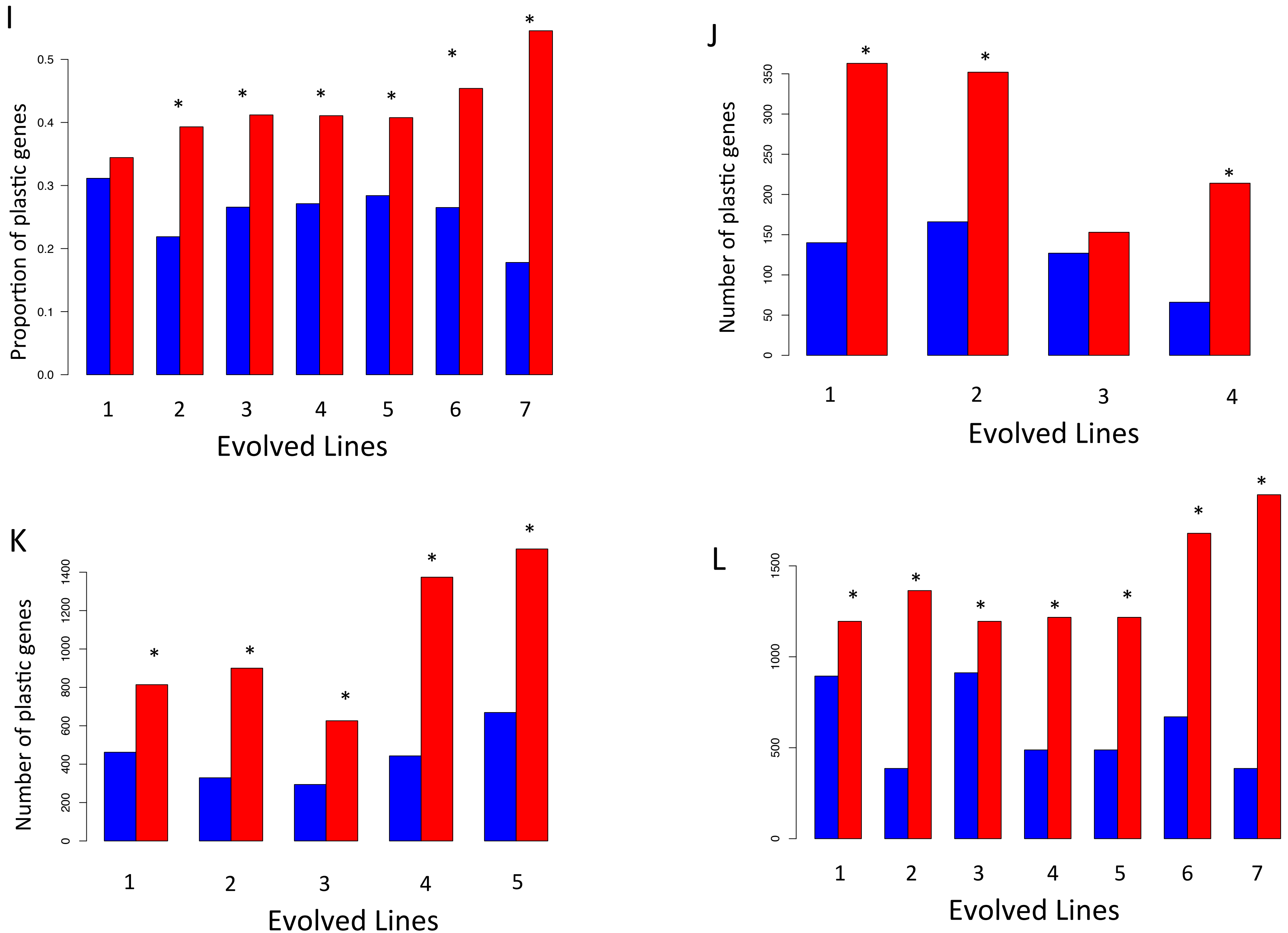
I
Proportion of plastic genes
0.0
0.1
0.2
0.3
0.4
0.5
1 2 3 4 5 6 7
Evolved Lines
J
Number of plastic genes
0
50
100
150
200
250
300
350
1 2 3 4
Evolved Lines
K
Number of plastic genes
0
200
400
600
800
1000
1200
1400
1 2 3 4 5
Evolved Lines
L
0
500
1000
1500
1 2 3 4 5 6 7
Evolved Lines

**Figure S3.1**: Proportion of genes showing a reversion or reinforcement of ancestral plasticity during evolution in different selection lines that adapted to dry, hot or hot-dry conditions. We tested the influence of different cutoffs for defining plastic and evolutionary responses. For **B, D, F** genes with plastic changes were defined as genes where absolute difference in expression levels between control and treatment conditions was higher than 50 % of ancestral levels. Genes with evolutionary changes were defined as genes where absolute difference of mean expression levels between selection lines and control line in the treatment was higher than 50 % of ancestral levels. Ancestral levels are expression levels of Control-lines in control conditions. **B**: Dry-lines; **D**: Hot-lines, **F**: Hot-Dry–lines. For **A, C, E** no cutoff was applied. **A**: Dry-lines, **C**: Hot-lines, **E**: Hot-Dry lines. **G** (Dry-lines),**H** (Hot-lines), **I** (Hot-Dry–lines) show results for a cutoff of 20%. This cutoff was used for analyses presented in the manuscript. Shown are proportions of plastic genes. **J, K, L** show results of a parametric bootstrap that was proposed by Ho and Zhang 2019 to test for prevalence of reversions. A gene was classified as showing a reversion (or reinforcement respectively) of the plastic response if at least 950 of 1000 repeats showed this pattern. **J:** Dry-lines, **K:** Hot-lines, **L**: Hot-Dry–lines. Differences in proportions were tested by a two-tailed binomial test. Significantly excess of reversions are indicated by *.

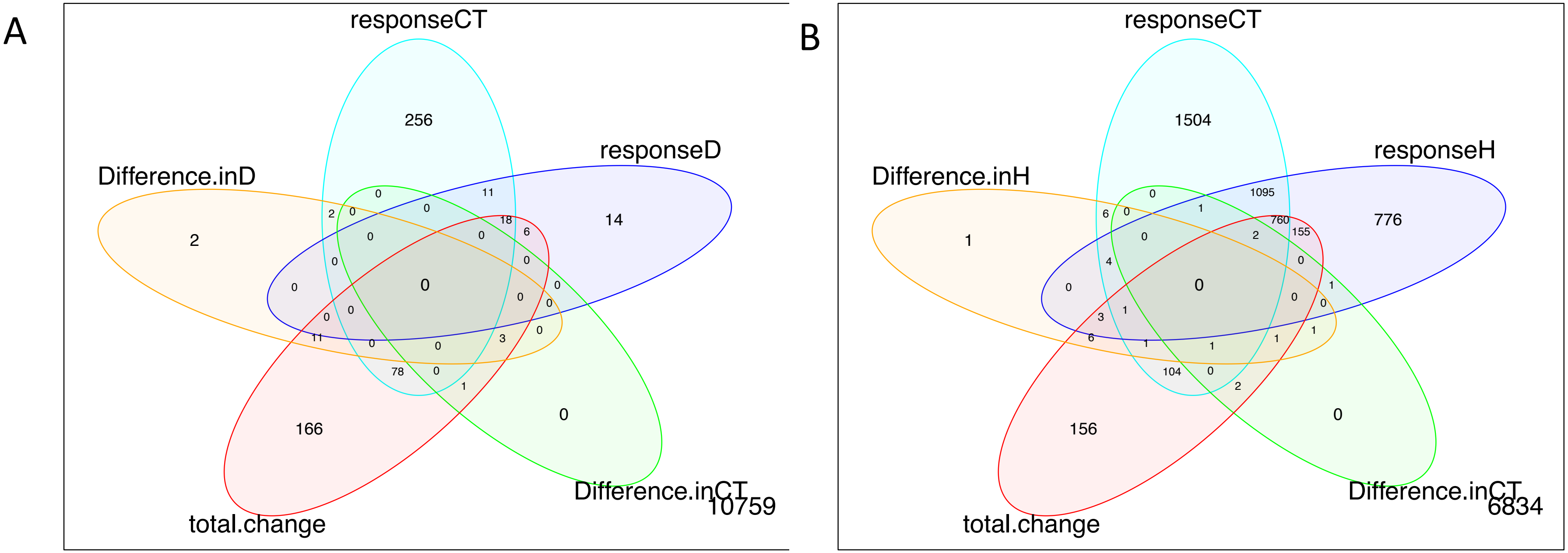


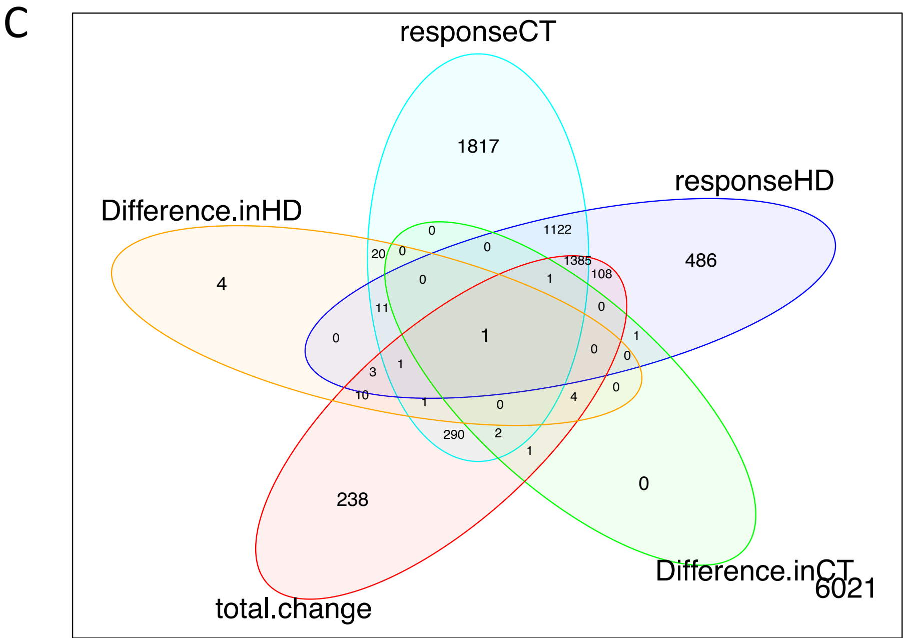


**Figure S3.2:** Number of differentially expressed (DE) genes in Dry **(A)**, Hot **(B)**, Hot-Dry **(C)**. responseCT: plastic response of Control-lines to treatment conditions; responseD, responseH, responseHD: plastic responses of selection lines; Difference in CT: differences between Control-lines and selection lines in control conditions; Difference in D, Difference in H, Difference in HD: differences in treatment conditions; total change: difference between Control-lines in control conditions and selection lines in their respective treatments. Intersections show genes that are DE in several comparisons.

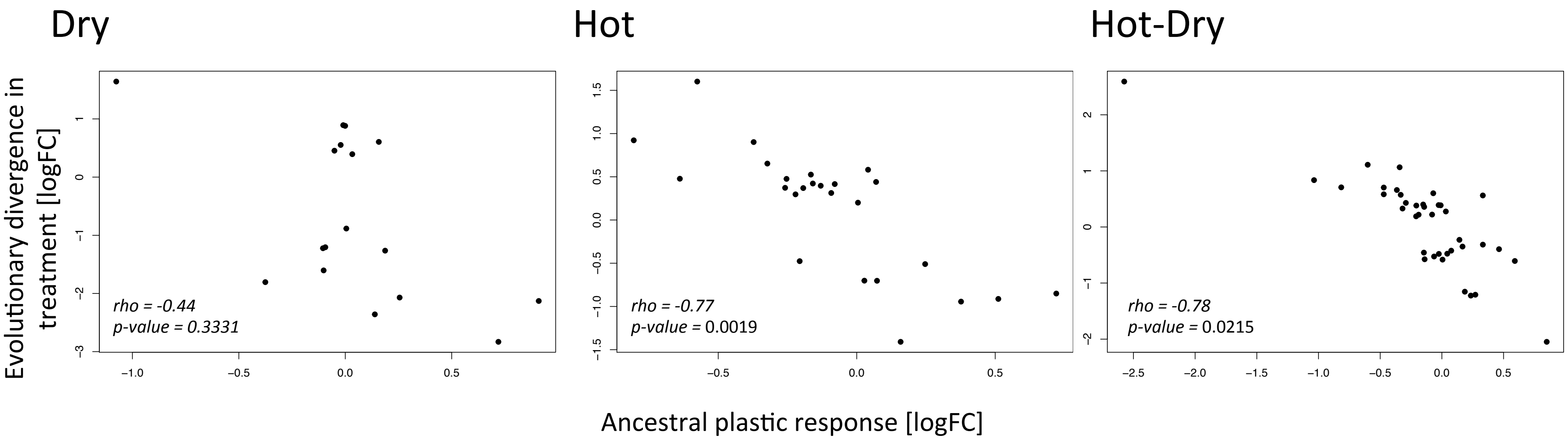


**Figure S3.3:** Correlation of ancestral plastic response (changes in gene expression in control-lines between treatment and control conditions) and evolutionary divergence in treatment (differences in gene expression levels between adapted selection lines and Control-lines. P-values were obtained from 10,000 permutations and give the proportion of permutations, which showed a more negative correlation than the observed one.
