## Supporting information S1 for "Restoring ancestral phenotypes by reduction of plasticity is a general pattern in gene expression during adaptation to different stressors in *Tribolium castaneum*"

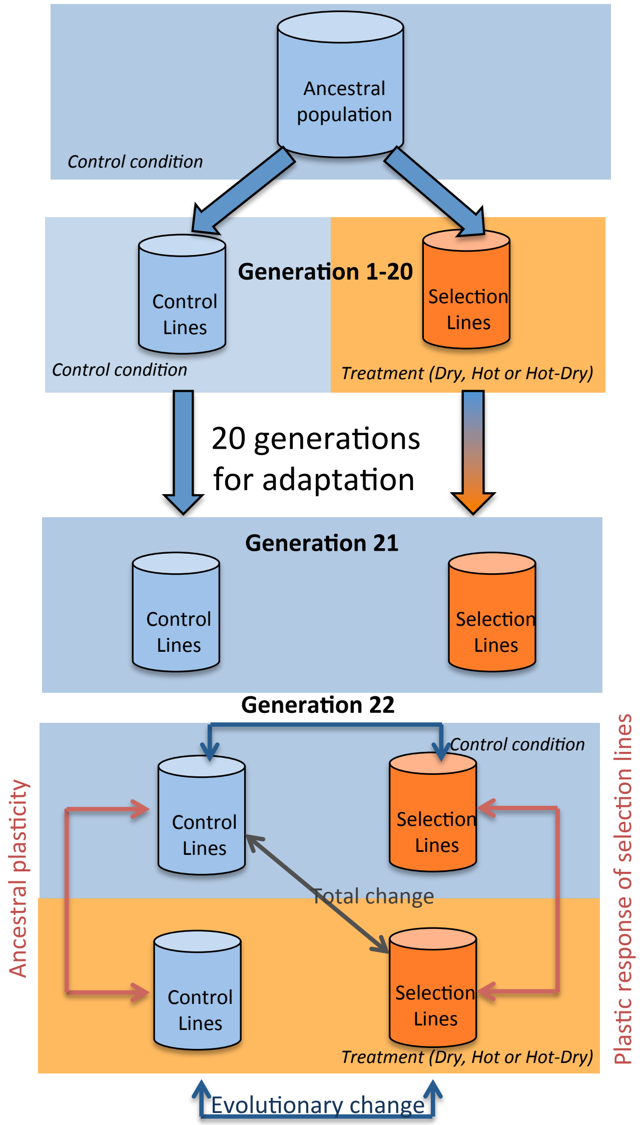


**Figure S1:** Schematic representation of the experimental evolution design. Lines were derived from a single outbred *Tribolium castaneum* population (collected in 2013, strain Cro1) that had lived under control conditions for ca. 30 generations. For each selection regime we used six replicate lines. In generation 15, four additional lines were created by mixing the six lines of each selection regime in equal proportions, resulting in a total of 10 lines per selection regime. Lines stayed in control or treatment conditions for 20 generations. After one generation in control condition to remove potential maternal and epigenetic effects, individuals from all lines were transferred to control and treatment conditions in the egg stage. Fitness and gene expression were measured in adult females. To get plastic responses in gene expression, expression levels in treatment conditions were compared to levels under control conditions of the same line. Evolved response is the difference within treatment conditions between control and selection lines. Control conditions: 33°C, 70% relative humidity r.h.; Treatments: Dry: 33°C, 30% r.h.; Hot 37°C, 70% r.h.; Hot-Dry: 37°C, 30% r.h.
