## Supporting information S2 for "Restoring ancestral phenotypes by reduction of plasticity is a general pattern in gene expression during adaptation to different stressors in *Tribolium castaneum*"

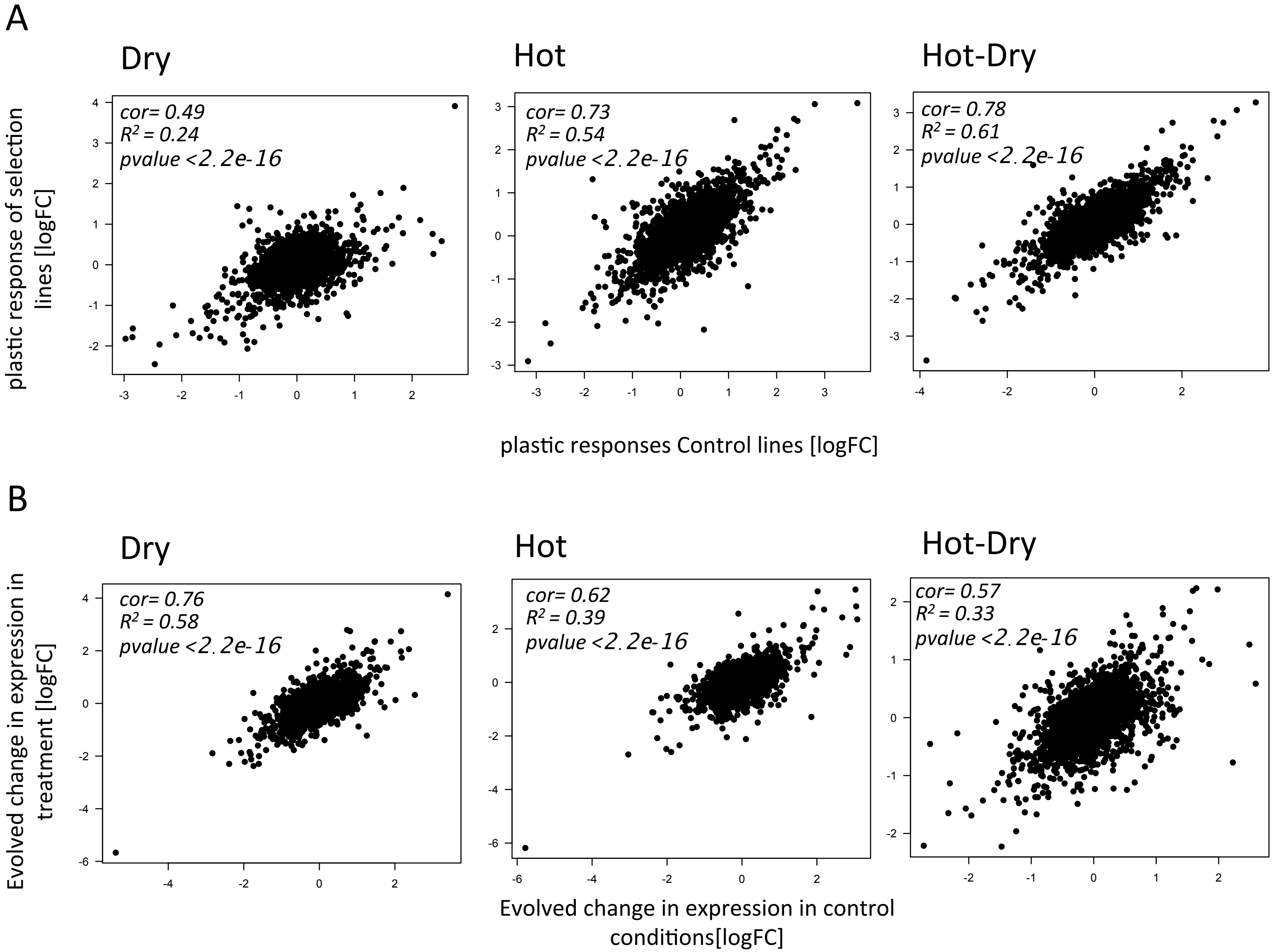


**Figure S2:** Correlation of plastic responses in gene expression of control lines and selection lines (**A**) and correlation of evolved divergence in gene expression in control and treatment conditions (**B**). Selection lines had lived in the respective treatment for 20 generations. After filtering lowly expressed genes (minimum of 1cpm in at least 2 samples) 11534 genes remained that were used for calculating correlations.
